# Maptcha: An efficient parallel workflow for hybrid genome scaffolding

**DOI:** 10.1101/2024.03.25.586701

**Authors:** Oieswarya Bhowmik, Tazin Rahman, Ananth Kalyanaraman

**Affiliations:** Washington State University, Pullman, WA, USA

**Keywords:** genome assembly, hybrid scaffolding, long read mapping, sketching

## Abstract

**Background:** Genome assembly, which involves reconstructing a target genome, relies on scaffolding methods to organize and link partially assembled fragments. The rapid evolution of long read sequencing technologies toward more accurate long reads, coupled with the continued use of short read technologies, has created a unique need for hybrid assembly workflows. The construction of accurate genomic scaffolds in hybrid workflows is complicated due to scale, sequencing technology diversity (e.g., short vs. long reads, contigs or partial assemblies), and repetitive regions within a target genome.

**Results:** In this paper, we present a new parallel workflow for hybrid genome scaffolding that would allow combining pre-constructed partial assemblies with newly sequenced long reads toward an improved assembly. More specifically, the workflow, called `Maptcha`, is aimed at generating long genome scaffolds of a target genome, from two sets of input sequences—an already constructed partial assembly of contigs, and a set of newly sequenced long reads. Our scaffolding approach internally uses an alignment-free mapping step to build a ⟨contig,contig⟩ graph using long reads as linking information. Subsequently, this graph is used to generate scaffolds. We present and evaluate a graph-theoretic “wiring” heuristic to perform this scaffolding step. To enable efficient workload management in a parallel setting, we use a batching technique that partitions the scaffolding tasks so that the more expensive alignment-based assembly step at the end can be efficiently parallelized. This step also allows the use of any standalone assembler of choice for generating the final scaffolds.

**Conclusions:** Our experiments with `Maptcha` on a variety of input genomes, and comparison against a state-of-the-art hybrid scaffolder (`LRScaf`) demonstrate that `Maptcha` is able to generate longer and more accurate scaffolds in significantly faster runtimes. For instance, `Maptcha` produces scaffolds with an NG50 length of 4.8Mbp (compared to 171Kbp by `LRScaf`) for *T. crassiceps*, and 81Mbp for Human chr 7 (compared to 4.5Mbp by `LRScaf`), while reducing the runtime from hours to minutes in several cases. We also performed a coverage experiment by varying the sequencing coverage depth for long reads, which demonstrated the potential of `Maptcha` to generate significantly longer scaffolds in low coverage settings (1× to 10×).

## Background

Advancements in sequencing technologies, and in particular, the continuous evolution from high throughput short read to long read technologies, have revolutionized biological sequence analysis. The first generation of long read technologies such as PacBio SMRT [27] and Oxford Nanopore Technologies (ONT) [10] sequencing platforms, were able to break the 10 Kbp barrier for read lengths. However, these technologies also carry a higher cost per base than short read (e.g., Illumina) platforms, and they also have much higher per-base error rate (5-15%). Recent long read technologies such as PacBio HiFi (High Fidelity) [43, 16, 40] have significantly improved accuracy (99.9%).

Genome assembly, irrespective of the sequencing approach employed, strives to accomplish three fundamental objectives. Firstly, it aims to reconstruct an entire target genome in as few pieces (i.e., contiguous sequences) as possible. Secondly, the goal is to ensure the highest accuracy at the base level. Lastly, the process seeks to minimize the utilization of computational resources. Short read assemblers effectively address the second and third objectives [6, 19, 42], while long read assemblers excel in achieving the first goal [21, 8].

In the realm of contemporary genome assembly, long read assemblers have adopted the Overlapping-Layout-Consensus (OLC) paradigm [21, 8, 20, 39, 33, 37] and de Bruijn graph approaches [44, 26, 38]. These assemblers utilize advanced algorithms that greatly accelerate the comparison of all-versus-all reads. Many long read assembly tools also perform error correction by representing long reads through condensed and specialized k-mers, such as minimizers [32] and minhashes [37]. This refined representation expedites the identification of overlaps exceeding 2 kb. The most recent long read assemblers are now progressing toward reducing computational resources. However, the assembly of uncorrected long reads introduces challenges, necessitating additional efforts in the form of consensus polishing [7, 23, 41]. Genome assembly polishing is a process aimed at enhancing the base accuracy of assembled contig sequences. Typically, long read assemblers undergo a singular round of long read polishing, followed by multiple rounds of polishing involving both long and short reads using third-party tools.

The rapid progress in sequencing technologies has provided researchers with extensive quantities of raw genomic data. However, the reconstruction of a complete and accurate genome from these fragmented sequences remains a challenging task due to inherent complexities, repetitive regions, and limitations of individual sequencing techniques. Genome assembly, the process of reconstructing these sequences into a coherent whole, heavily relies on scaffolding methods to arrange and link these fragments.

Scaffolding is a process that bridges these technological gaps by harnessing additional information beyond the raw sequence data. Scaffolding acts as the architectural backbone, utilizing paired-end information, mate-pair relationships, or long-range linking data to order, orient, and link contigs, thereby stitching together the genomic fragments into a more accurate representation of the original genome. However, the reliance on a single sequencing technology for scaffolding could result in incomplete or fragmented assemblies [2]. This limitation necessitates hybrid scaffolding approaches that integrate multiple sequencing technologies and complementary data types. By combining the strengths of short read and long read sequencing along with other supplementary data such as optical mapping or Hi-C, hybrid scaffolding aims to overcome the deficiencies of individual methods and achieve more contiguous and accurate genome reconstructions.

### Hybrid scaffolding

The integration of contigs and long read information for scaffolding purposes can be a promising approach to improve existing genome assemblies [25]. Assemblies generated from short reads are known for their high accuracy, but are often limited by shorter contig lengths, as measured by N50 or NG50. On the other hand, long read sequencing technologies can span larger sections of the genome but are often hindered by higher costs which limit their sequencing coverage depths (to typically under 20× vs. 100× for short reads), and higher error rates compared to short read sequencing complicating *de novo* assembly. Hybrid scaffolding workflows can overcome these limitations by integrating the fragmented assemblies of contigs obtained from short reads and utilizing the long reads to order and orient contigs into longer scaffolds.

In this paper, we visit the *hybrid scaffolding* problem. Given an input set of contigs (C) generated from short reads, and a set of long reads (ℒ), hybrid scaffolders aim to order and orient the contigs in C using linking information inferred from the long reads ℒ. Such an approach has the advantage of reusing and building on existing assemblies to create improved versions of assemblies incrementally, as more and more long read sequencing data sets are available for a target genome.

### Related work

While the treatment of the hybrid scaffolding problem is more recent, there are several tools that incorporate long read information for extending contigs into scaffolds. The concept of genome scaffolding initially emerged in the realm of classical *de novo* genome assembly, as introduced by Huson et al. [18]. This pioneering work aimed to arrange and align contigs utilizing paired-end read information alongside inferred distance constraints. Of the two steps in scaffolding, the alignment step is not only computationally expensive, but it can also lead to loss in recall using traditional mapping techniques. On the other hand, the second step of detecting the true linking information between contig pairs can be prone to false merges, impacting precision—particularly for repetitive genomes.

Over subsequent years, a suite of tools emerged within this classical framework, each striving to refine scaffolding methodologies [2, 9, 12, 13, 24, 28, 34, 35]. For an exhaustive exploration of these methods, refer to the comprehensive review by Luo et al. [25]. Most of these tools utilize alignments of long reads to the contigs of a draft assembly to infer joins between the contig sequences. The alignment information is subsequently used to link pairs of contigs that form successive regions of a scaffold. SSPACE-LongRead produces final scaffolds in a single iteration and has shown to be faster than some of the other scaffolders for small eukaryaotic genomes; but it takes very long runtimes on larger genomes. For instance, SSPACE-LongRead takes more than 475 hours to assemble *Z*.*mays* and for the *Human CHR*1, it takes more than a month. OPERA-LG [14] provides an exact algorithm for large and repeat-rich genomes. It requires significant mate-pair information to constrain the scaffold graph and yield an optimised result. OPERA-LG is not directly designed for the PacBio and ONT data. To construct scaffold edges and link contigs into scaffolds, OPERA-LG needs to simulate and group mate-pair relationship information from long reads.

`LRScaf` [29] is one of the most recent long read scaffolding tools which utilizes alignment tools like BLASR [4] or `Minimap2` [22] to align contigs with long reads, and generates alignment information. These alignments form the basis for establishing links between contigs. Subsequently, a scaffold graph is constructed, wherein vertices represent contig ends, and edges signify connections between these ends and associated long reads. This graph also encapsulates information regarding contig orientation and long read identifiers. To mitigate errors and complexities arising from repeated regions and high error rates, `LRScaf` meticulously refines the scaffold graph. This refinement process involves retaining edges associated with a minimal number of long reads and ensuring the exclusion of edges connecting nodes already present within the graph. The subsequent stage involves navigating linear stretches of the scaffold graph. `LRScaf` traverses the graph, systematically identifying linear paths until encountering a divergent node, signifying a branching point. At this juncture, the traversal direction is reversed, ensuring exhaustive exploration of unvisited and distinct nodes within the graph. This iterative process continues until all unique nodes are visited, resulting in a complete set of scaffolds from the linear paths within the graph. As can be expected, this rigorous process can be time-consuming, taking hours of compute time even on medium sized genomes (as shown later in the Results).

### Contributions

We present a new scalable algorithmic workflow, called `Maptcha`, for carrying out hybrid scaffolding on parallel machines, on inputs which are contigs (C) and high fidelity long reads (L). Figure 1 illustrates the major phases of the `Maptcha` workflow. Our approach is a graph-theoretic approach that first constructs a contig graph based on the mapping information between long reads and contigs, and subsequently uses that graph to generate the scaffolds. More specifically, the approach uses the following key ideas: a) use of a sketching-based alignment-free *mapping* step to build and refine the graph progressively between contig expansion and long read island construction; b) a vertex-centric heuristic called *wiring* to generate ordered walks of contigs from the graph that correspond to partial scaffolds; and c) a final *linking* step to bridge the partial scaffolds to create the final set of scaffolds.

**Figure 1.**
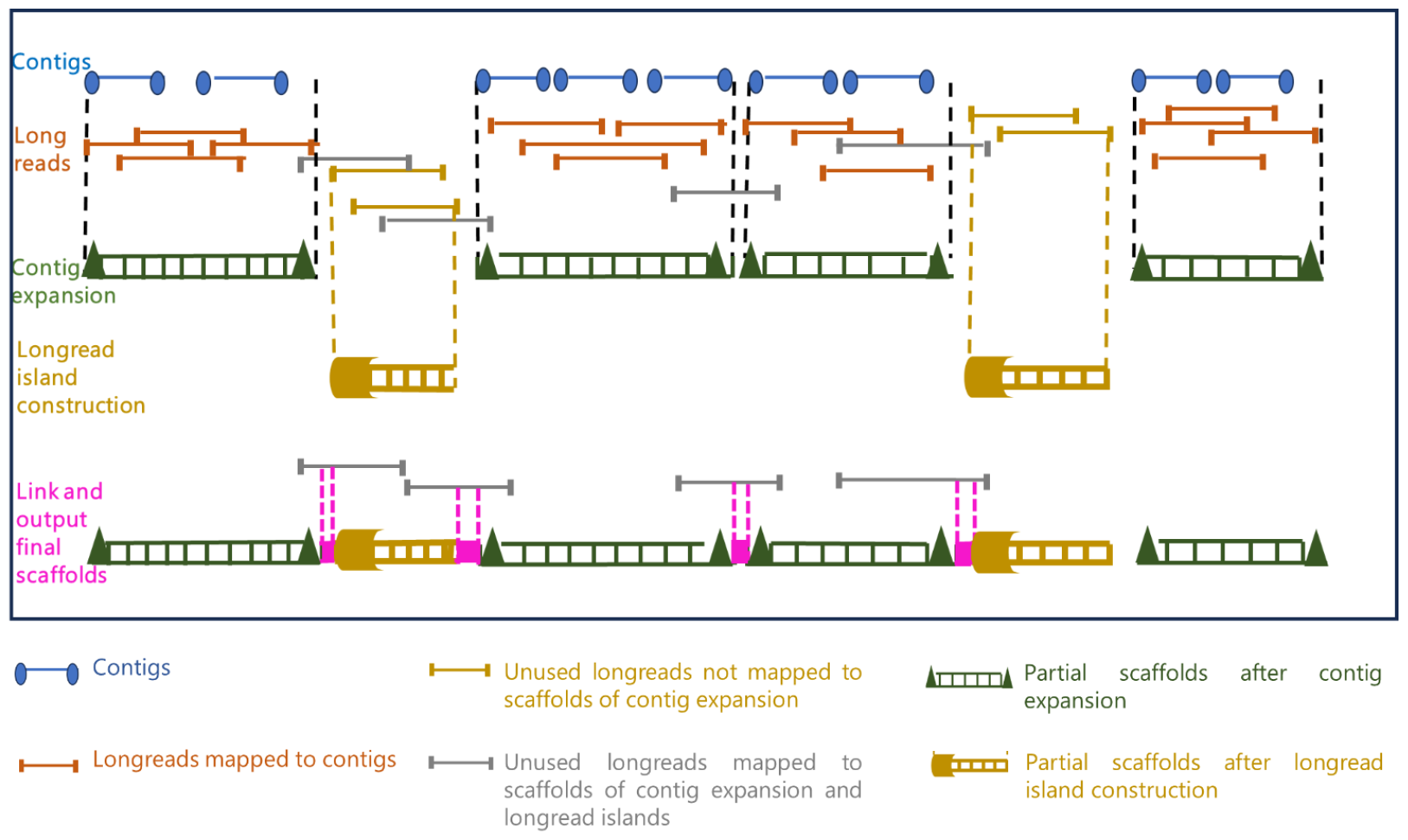
A schematic illustration of the major phases of the proposed Maptcha approach.

To enhance scalability, we have implemented a parallel batching technique for scaffold generation, which enables any standalone assembler to run in a distributed parallel manner while generating high-quality scaffolds. We use `Hifiasm` [5] as the standalone assembler for this paper, and use `JEM-mapper` [30, 31] for implementing the mapping step.

Our experiments with `Maptcha` on a variety of input genomes, suggest that `Maptcha` is able to generate longer and more accurate scaffolds than the state-of-the-art hybrid scaffolder `LRScaf`, while drastically reducing time to solution. For example, the scaffolds produced on the test input *D. busckii*, yielded an NG50 of 3,533 Kbp without any misassemblies (compared to an NG50 of 2,597 Kbp with 42 misassemblies for `LRScaf`). When handling a larger input data set such as *H. aestivaria*, `Maptcha` also demonstrated significant speedup, by completing in 1,841 seconds, while `LRScaf` could not complete within 6 hours.

We also compared `Maptcha` with a standalone state-of-the-art long read assembler running only on long reads (`Hifiasm (only-LR)`) to demonstrate the value of adding contigs to long read sets. Results showed that `Maptcha` is able to improve the quality of the overall assembly (i.e., longer scaffolds and reduced misassemblies) and faster runtimes. Notably, we also performed a coverage experiment by varying the sequencing coverage depth for long reads, which demonstrated the potential of `Maptcha` to generate significantly longer scaffolds in low coverage settings (1× to 10×).

The `Maptcha` software is available as open source for download and testing at https://github.com/Oieswarya/Maptcha.git.

## Methods

In this section, we describe in detail all the steps of our `Maptcha` algorithmic framework for hybrid scaffolding. Let *𝒞* = {*c*_1_, *c*_2_, … *c*_*n*_} denote a set of *n* input contigs (from prior assemblies). Let *ℒ* = {*r*_1_, *r*_2_, … *r*_*m*_} denote a set of *m* input long reads. Let |*s*| denote length of any string *s*. We use 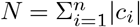 and 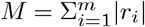. Furthermore, for contig *c*, let 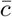 denote its reverse complement.

### Problem statement

Given *𝒞* and *ℒ*, the goal of our hybrid scaffolding problem is to generate a set of scaffolds *𝒮* such that a) each scaffold *S* ∈ *𝒮* represents a subset of *𝒞* such that no two subsets intersect (i.e., *S*_*i*_ ∩ *S*_*j*_ = ∅); and b) each scaffold *S* ∈ *𝒮* is an ordered sequence of contigs [*c*_1_, *c*_2_, …], with each contig participating in either its direct form *c* or its reverse complemented form 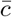. Here, each successive pair of contigs in a scaffold is expected to be linked by one or more long reads *r* ∈ *ℒ*. Intuitively, there are two objectives: i) maximize recall—i.e., to generate as few scaffolds as possible, *and* ii) maximize precision—i.e.,the relative ordering and orientation of the contigs within each scaffold matches the true (but unknown) ordering and orientation of those contigs along the target genome.

### Algorithm

The design of the `Maptcha` scaffolding algorithmic framework is broken down into three major phases.

- `contig expansion`: In the first phase, using the contigs as seeds, we aim to extend them on either end using long reads that align with those contigs. This extension step is also designed to detect and connect successive pairs of contigs with direct long read links. This yields the first generation of our partial scaffolds.
- `longread island construction`: Note that not all long reads may have contributed to these partial scaffolds, in particular those long reads which fall in the gap regions of the target genome between successive scaffolds. Therefore, in the next phase, we detect the long reads that do not map to any of the first generation partial scaffolds, and use them to build partial scaffolds corresponding to these long read island regions. This new set of partial scaffolds corresponds to the second generation of partial scaffolds.
- `link scaffolds with bridges`: Finally, in the last phase, we aim to link the first and second generation scaffolds using long reads that serve as bridges between them. This step outputs the final set of scaffolds.

This three phase approach has the following **advantages**. First, it provides a systematic way to progressively combine the sequence information available from the input contigs (which typically tend to be more accurate albeit fragmented, if generated from short reads) to the input long reads (which may be significantly larger in number), in an incremental fashion. Next, this incremental approach also could reduce the main computational workload within each phase that is required for mapping long reads. More specifically, we choose to align long reads either to the contigs or to the generated partial scaffolds wherever possible, and in the process restrict the more time consuming long read to long read alignments only to the gap regions not covered by any of the contigs or partial scaffolds. Additionally, our framework is capable of leveraging any off-the-shelf long read mapping tool. In this paper, we use the `JEM-mapper`, which is a recently developed fast (parallel) and accurate sketch-based alignment-free long read mapping tool suited for hybrid settings [30, 28]. Finally, by decoupling the contig ordering and orientation step (which is a graph-theoretic problem) from the scaffold generation step (which is an assembly problem), we are able to efficiently parallelize the scaffold generation step. This is achieved through a batching step that splits the input sequences into separate batches to allow the use of any existing standalone long read assembler to generate the final sequence scaffolds. In this paper, we use `Hifiasm` [5], which is one of the most widely used state-of-the-art long read assembly tool, as our standalone assembler.

In what follows, we describe the details of our algorithm for each of the three major phases of our approach. Figure 2 provides an illustration of all the main steps within each of these three phases.

**Figure 2.**
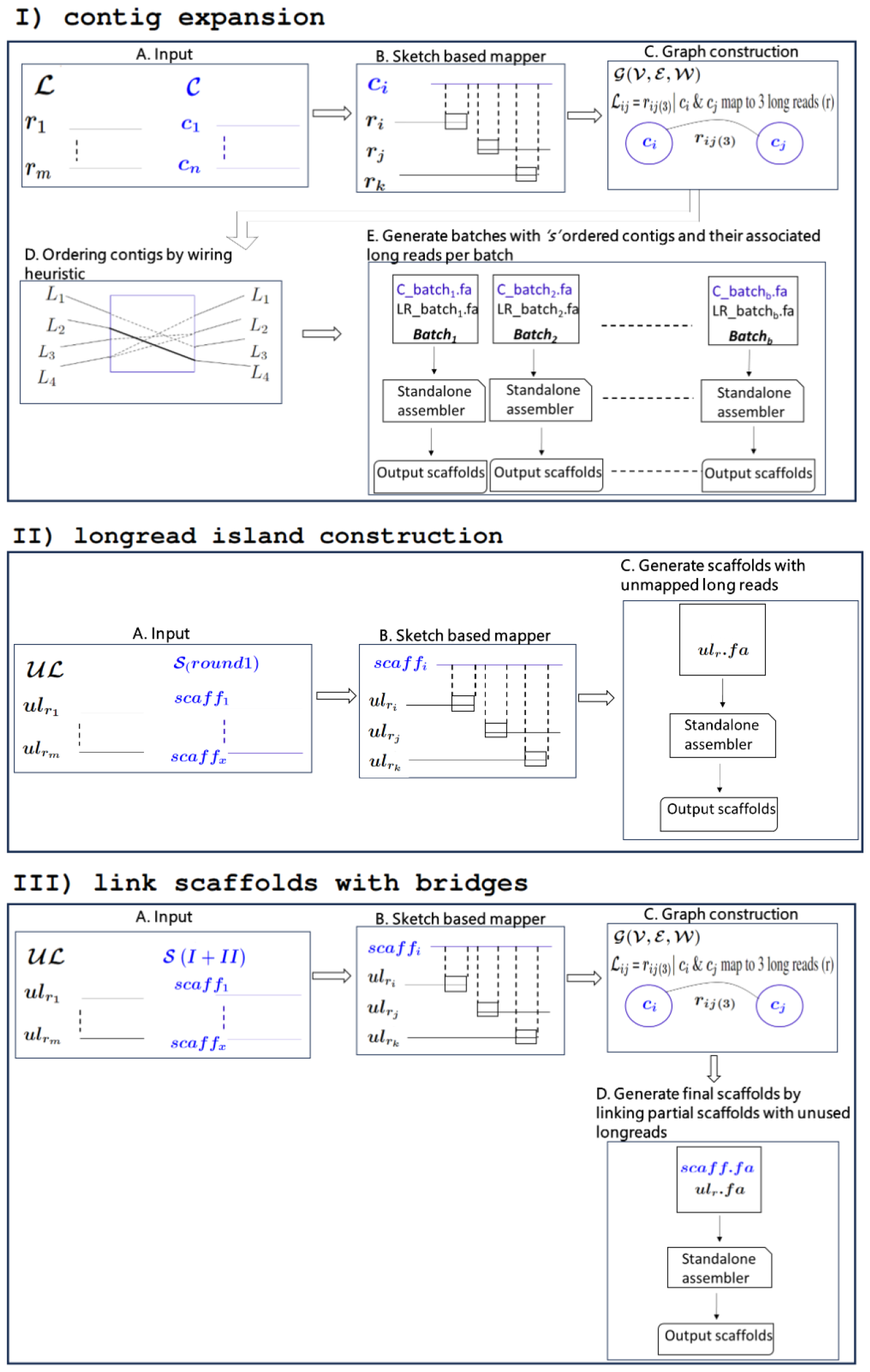
A detailed illustration of the Maptcha pipeline showing the different phases and their steps.

### Phase: contig expansion

The goal of this phase is to enhance contigs by incorporating long reads that have been aligned with them. This process allows for the extension of contigs by connecting multiple ones into a scaffold using the long reads aligned to them, thereby increasing the overall length of the contigs. This is achieved by first mapping the long reads to contigs to detect those long reads that map to contigs, and then use that information to link contigs and extend them into our first generation of partial scaffolds (panel I in Figure 2).

We use the following definition of a partial scaffold in our algorithm: A *partial scaffold* corresponds to an ordered and oriented sequence of an arbitrary number of contigs [*c*_*i*_, *c*_*j*_, *c*_*k*_, …] such that every consecutive pair of contigs along the sequence are linked by one or more long reads.

#### Step: Mapping long reads to contigs

For mapping, we use an alignment-free, distributed memory parallel mapping tool, `JEM-mapper` because it is both fast and accurate [30, 31]. `JEM-mapper` employs a sketch-based alignment-free approach that computes a minimizer-based Jaccard estimator (`JEM`) sketch between a subject sequence and a query sequence. More specifically, in a preprocessing step, the algorithm generates a list of minimizing *k*-mers [32, 36] from each subject (i.e., each contig) and then from that list computes minhash sketches [3] over *T* random trials (we use *T* = 30 for our experiments). Subsequently, `JEM` sketches are generated from query long reads. Based on these sketches, for each query the tool reports the subject to whom it is most similar. For further details on the methodology, refer to the original paper by Rahman et al. [30].

One challenge of using a mapping tool is that the subject (contigs) and query (long reads) sequences may be of variable lengths, thereby resulting possibly in vastly different sized ground sets of minimizers from which to draw the sketches. However, it is the minimizers from the *aligning region* between the subject and query that should be ideally considered for mapping purposes. To circumvent this challenge, in our implementation we generate sketches only from the two ends of a long read. In our implementation, we used a length of *𝓁* base pairs (*𝓁* = 2*Kbp* used in our experiments) from either end of a long read for this purpose. The intuitive rationale is that since we are interested in a scaffolding application, this approach of involving the ends of long reads (and their respective alignment with contigs) provides a way to link two distantly located contigs (along the genome) through long read bridges.

Using this approach in our preliminary experiments, we compared `JEM-mapper` with `Minimap2` and found that `JEM-mapper` yielded better quality results for our test inputs (results summarized in the supplementary section Figure S2).

#### Step: Graph construction

Note that the mapping step maps every long read to at most one contig. However, a contig can be potentially mapped to multiple long reads and the number of such long reads mapped to a contig is dependent on the sequencing coverage depth. Let *ℳ* denote the mapping output, which can be expressed as the set of 2-tuples of the form ⟨*c, r*⟩—where long read *r* maps to a contig *c*—output by the mapper. We use *L*_*c*_ ⊆ *ℒ* to denote the set of all long reads that map to contig *c*, i.e., *L*_*c*_ = {*r* | ⟨*c, r*⟩ ∈ *ℳ*}. Informally, we refer to *L*_*c*_ as the *long read set corresponding to contig c*.

Using the information in *ℳ*, and in *L*_*c*_ for all *c* ∈ *𝒞*, we construct an undirected graph *G*(*V, E*), where:

- *V* is the vertex set such that there is one vertex for every contig *c* ∈ *𝒞*; and
- *E* is the set of all edges of the form (*e*_*i*_, *e*_*j*_), such that there exists at least one long read *r* that maps to both contigs *c*_*i*_ and *c*_*j*_ (i.e., 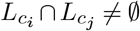).

Intuitively, each edge is the result of two contigs sharing one or more long reads in their mapping sets. In our implementation, we store the set of long read IDs corresponding to each edge. More specifically, along with an edge (*e*_*i*_, *e*_*j*_) ∈ *E*, we also store its long read set *L*_*i,j*_ given by the set 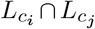. The cardinality of set *L*_*i,j*_ is referred to as the “support value” for the edge between these two contigs. Since the vertices of *G* correspond to contigs, we refer to *G* as a *contig graph*.

Next, the graph *G* along with all of its auxiliary edge information as described above, are used to generate partial scaffolds. We perform this in two steps: a) First enumerate paths in the contig graph that are likely to correspond to different partial scaffolds (this is achieved by our wiring algorithm that is described next); and b) subsequently, generate contiguous assembled sequences for each partial scaffold by traversing the paths from the previous step (this is achieved by using a batch assembly step described subsequently).

#### Step: Wiring heuristic

Recall that our goal is to enumerate partial scaffolds, where each partial scaffold is a maximal sequence of contiguously placed (non-overlapping) contigs along the target genome. In order to enumerate this set of partial scaffolds, we make the following observation about paths generated from the contig graph *G*(*V, E*). A partial scaffold [*c*_*i*_, *c*_*i*+1_, …, *c*_*j*_] can be expected to be represented in the form of a path in *G*(*V, E*). However, it is important to note that not all graph paths may correspond to a partial scaffold. For instance, consider a branching scenario where a path has to go through a branching node where there are more than one viable path out of that node (contig). If a wrong decision is taken to form paths out of branching nodes, the resulting paths could end up having chimeric merges (where contigs from unrelated parts of the genome are collapsed into one scaffold). While there is no way to check during assembly for such correctness, we present a technique we call *wiring*, as described below, to compute partial scaffolds that reduce the chance of false merges.

The wiring algorithm’s objective is one of enumerating maximal acyclic paths in *G*—i.e., maximality to ensure longest possible extension of the output scaffolds, and acyclic to reduce the chance of generating chimeric errors due to repetitive regions in the genome (as illustrated in Figure 5). This problem is trivial if each vertex in *V* has at most two neighbors in *G*, as it becomes akin to a linked list of contigs, each with one *predecessor* contig and one *successor* contig. However, in practice, we expect several branching vertices that have a degree of more than two (indicating potential presence of repeats). Therefore, finding a successor and/or a predecessor vertex becomes one of a non-trivial path enumeration problem that carefully resolves around branching nodes.

### Algorithm

Our wiring algorithm is a linear time algorithm that first computes a “wiring” internal to each vertex, between edges incident on each vertex, and then uses that wired information to generate paths. First, we described the wiring heuristic.

Step 1: Wiring of vertices

For each vertex *c* ∈ *V* that has at least degree two, the algorithm selects a subset of two edges incident on that vertex to be “wired”, i.e., to be connected to form a path through that vertex, as shown in Figure 3. The two edges so wired determine the vertices adjacent on either side of the current vertex *c*.

**Figure 3.**
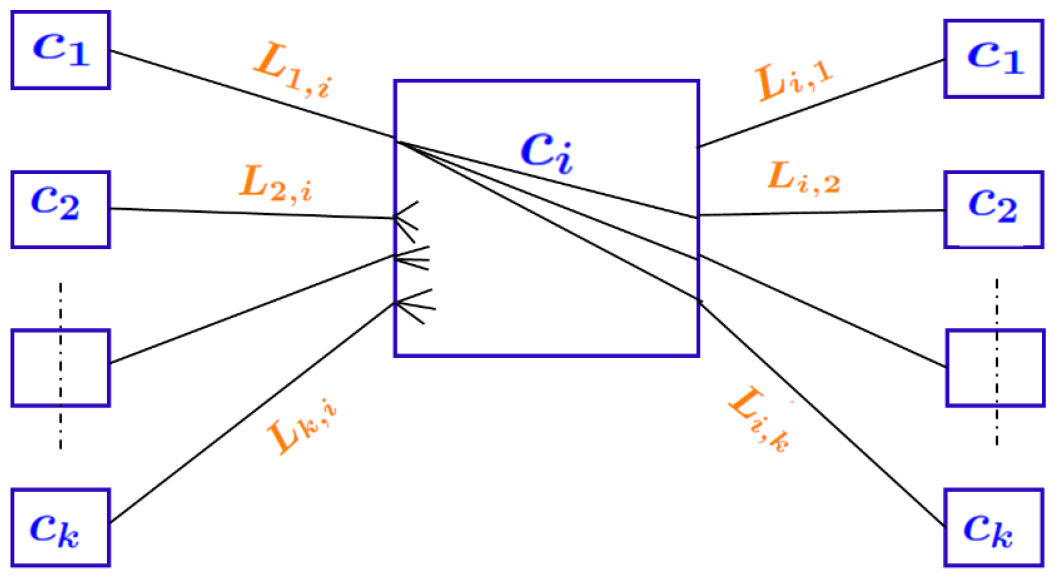
Illustration of the wiring heuristic, shown at a contig vertex *c*_*i*_. Out of all possible pairwise connections between the incident edges, only one edge pair is chosen—i.e., “wired”.

To determine which pair of edges to connect, we use the following heuristic. Let *L*_*i*_ denote the set of long read IDs associated with edge *e*_*i*_. We then *(hard) wire* two edges *e*_*i*_ and *e*_*j*_ (*c*_*i*_ ≠ *c*_*j*_) at vertex *c*, if *L*_*i*_ ∩ *L*_*j*_ ≠ *ϕ* and it is maximized over all possible pairs of edges incident on *c*, i.e., 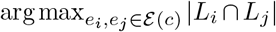, where E(*c*) denotes all edges incident on *c*. The simple intuition is to look for a pair of edges that allows maximum long read-based connectivity in the path flowing through that vertex (contig). This path has the largest *support* by the long read set and is therefore most likely to stay true to the connectivity between contigs along the target genome. All other possible paths through that vertex are ignored. The resulting wired pair of edges ⟨*e*_*i*_, *e*_*j*_⟩ generated from each vertex *c* is added in the form of wired edge 3-tuple ⟨*c*_*i*_, *c*_*j*_, *c*⟩. We denote the resulting set as *𝒲*.

As a special case, if a vertex *c* has degree one, then the wiring task is trivial as there exists only one choice to extend a path out of that contig, *c*_*e*_, along the edge *e* attached to that vertex. We treat this as a special case of wiring by introducing a dummy contig *c*_*d*_ to each such vertex with degree one, and adding the tuple ⟨*c*_*d*_, *c*_*e*_, *c*⟩ to *𝒲*.

Note that by this procedure, each vertex *c* has at most one entry in *𝒲*. To implement this wiring algorithm, note that all we need is to store the set of long read *IDs* along each edge. A further advantage of this approach is that this is an independent decision made at each vertex, and therefore this step easily parallelizes into a distributed algorithm that works with a partitioning of the input graph.

Step 2: Path enumeration

In the next step, we enumerate edge-disjoint acyclic paths using all the wired information from *𝒲*. The rationale behind the edge-disjoint property is to reduce the chance of genomic duplication in the output scaffolds. The rationale for avoiding cycles in paths is two-fold—both to reduce genomic duplication due to repetitive contigs, as well as to reduce the chance of creating chimeric scaffolds.

The path enumeration algorithm (illustrated through an example in Figure 4) works as follows.

**Figure 4.**
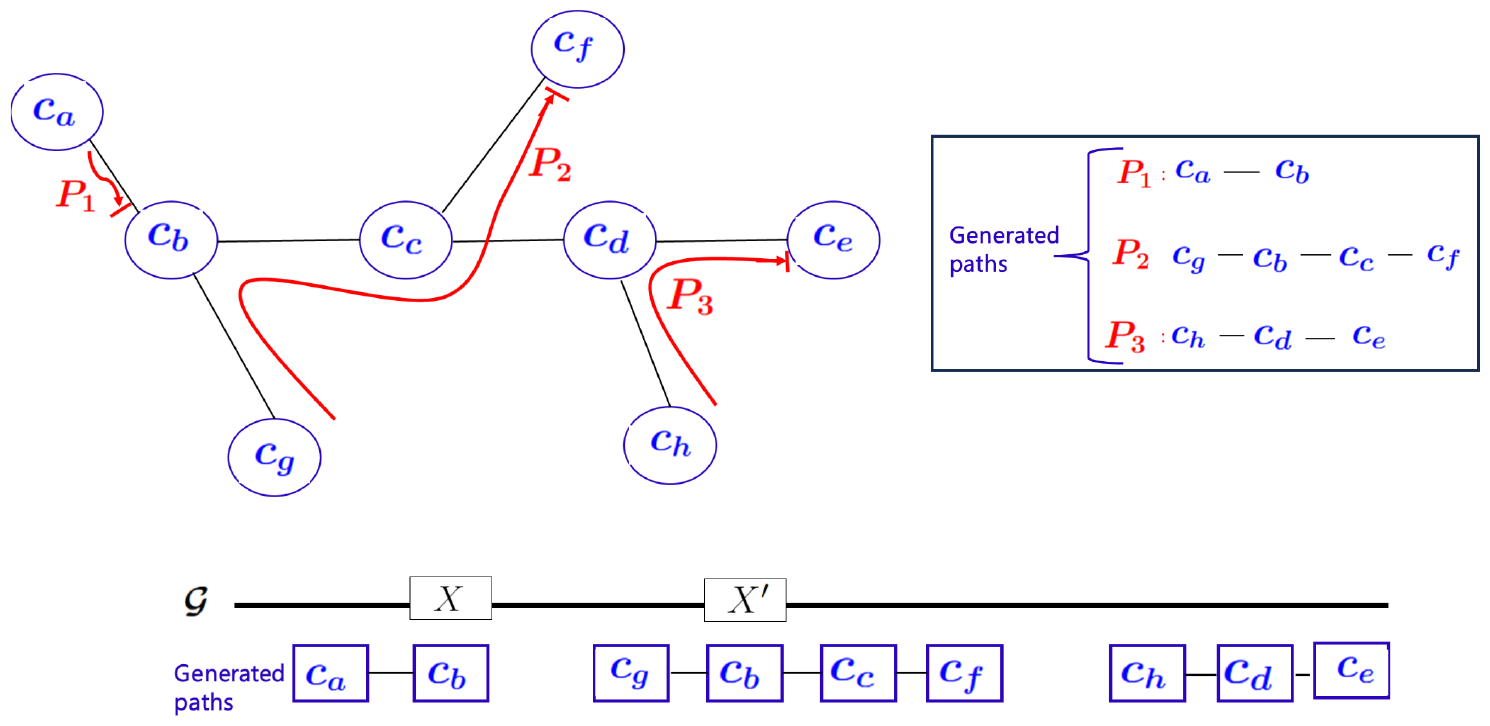
Edge-disjoint acyclic paths generated from walking the *contig-contig* graph. Also shown below are the likely alignments of the individual paths to the (unknown) target genome *𝒢*. Here, since the contig *c*_*b*_ appears in two paths, it is likely to be contained in a repetitive region (*X, X*^*′*^) as highlighted.

i. Initialize a *visit* flag at all vertices and set them to *unvisited*.
ii. Initialize a work queue *Q* of all vertices with degree one (e.g., *c*_*a*_, *c*_*e*_, *c*_*f*_, *c*_*g*_ and *c*_*h*_ in Figure 4).
iii. For each vertex *c* ∈ *Q*, if *c* is still unvisited, dequeue *c*, start a new path at *c* (denoted by *P*_*c*_), and grow the path as follows. The edge *e* incident on *c* connects *c* to another vertex, say *c*_1_. Then *c*_1_ is said to be the *successor* of *c* in the path and is appended to *P*_*c*_. We now mark the vertex *c* as visited. Subsequently, the algorithm iteratively extends the path by simply following the wired pairing of edges at each vertex visited along the way—marking each such vertex as visited and stitching together the path—until we arrive at one of the following termination conditions: Case a) Arrive at a vertex which has chosen a different predecessor vertex: See for example path *P*_1_ truncated at *c*_*b*_ because the wiring at *c*_*b*_ has chosen a different pair of neighbors other than *c*_*a*_ based on long read support, i.e., *𝒲* contains ⟨*c*_*g*_, *c*_*c*_, *c*_*b*_⟩. In this case, we add the vertex *c*_*b*_ at the end of the current path *P*_1_ and terminate that path. Case b) Arrive at a vertex that is already visited: This again implies that no extension beyond this vertex is possible without causing duplication between paths, and so the case is handled the same way as Case *a* by adding the visited vertex as the last vertex in the path and the path terminated. Case c) Arrive at a degree one vertex: This implies that the path has reached its end at the corresponding degree one contig and the path is terminated at this contig. More examples of paths are shown in Figure 4.

### Provable properties of the algorithm

The above wiring and path enumeration algorithms have several key properties.

#### Prop1

**Edge disjoint paths:** *No two paths enumerated by the wiring algorithm can intersect in edges. Proof:* This is guaranteed by the wiring algorithm (step 1), where each vertex chooses only two of its incident edges to be wired to build a path. More formally, by contradiction let us assume there exists an edge *e* that is covered by two distinct paths *P*_1_ and *P*_2_. Then this would imply that both paths have to pass through at least one branching vertex *c* such that there exist ⟨*e*_1_, *e, c*⟩ ∈ *𝒲* and ⟨*e*_2_, *e, c*⟩ ∈ *𝒲* (for some *e*_1_ ≠ *e*_2_ ≠ *e* all incident on *c*). However, by construction of the wiring algorithm (step 1) this is *not* possible. ▄

#### Prop2

**Acyclic paths:** *There can be no cycles in any of the paths enumerated. Proof:* This is guaranteed by the path enumeration algorithm described above (step 2). More specifically, the termination conditions represented by the Cases (a) and (b) clip any path before it forms a cycle. ▄

#### Prop3

**Deterministic routing:** *The path enumeration algorithm is deterministic and generates the same output set of paths for a given* W *regardless of the order in which paths are generated. Proof:* This result follows from the fact that the wiring heuristic at each vertex is itself deterministic as well as by the conditions represented by Cases (a) and (b) to terminate a path in the path enumeration algorithm. More specifically, note that each vertex contributes at most one hard-wired edge pair into *𝒲* and none of the other edge pair combinations incident on that vertex could lead to paths. Given this, consider the example shown in Figure 4, of two paths *P*_1_ and *P*_2_ converging onto vertex *c*_*b*_. Note that in this example, ⟨*c*_*g*_, *c*_*c*_, *c*_*b*_⟩ ∈ *𝒲*. The question here is if it matters whether we start enumerating *P*_1_ first or *P*_2_ first. The answer is no. In particular, if *P*_1_ is the first to get enumerated, then termination condition Case (a) would apply to terminate the path to end at *c*_*b*_. Therefore, when *P*_2_ starts, it will still be able to go through *c*_*b*_. On the other hand, if *P*_2_ is the first path to get enumerated, then *c*_*b*_ will get visited and therefore termination condition Case (b) would apply to terminate *P*_1_ at *c*_*b*_ again. So either way, the output paths are the same. A more detailed example for this order agnostic behavior is shown in S3. This order agnostic property allows us to parallelize the path enumeration process without having to synchronize among paths. ▄

As a corollary to Prop1 (on edge disjoint paths) and Prop2 (on acyclic paths), we now show an important property about the contigs from repetitive regions of the genome and how the wiring algorithm handles those contigs carefully so as to reduce the chances of generating chimeric scaffolds.

### Corollary 1

Let *c*_*x*_ be a contig that is completely contained within a repetitive region. Then this contig can appear as a non-terminal vertex^1^ in at most one path output by the wiring algorithm.

*Proof:* Consider the illustrative example in Figure 5, which shows a contig *c*_*x*_ that maps to a repeat *X* and its copy *X*^*′*^. In particular, if there is a trail of long reads linking the two repeat copies (from [*c*_*x*_, *c*_2_, … *c*_*k*_*′, c*_*x*_]), then it could generate a cycle in the graph *G*. However, based on Prop2, the cycle is broken by the path enumeration algorithm and therefore *c*_*x*_ is allowed to appear as a non-terminal vertex only in at most one of the paths that goes through it. Even if there is no trail of long reads connecting the two repeat regions, the same result holds because of the edge disjoint property of Prop1.

**Figure 5.**
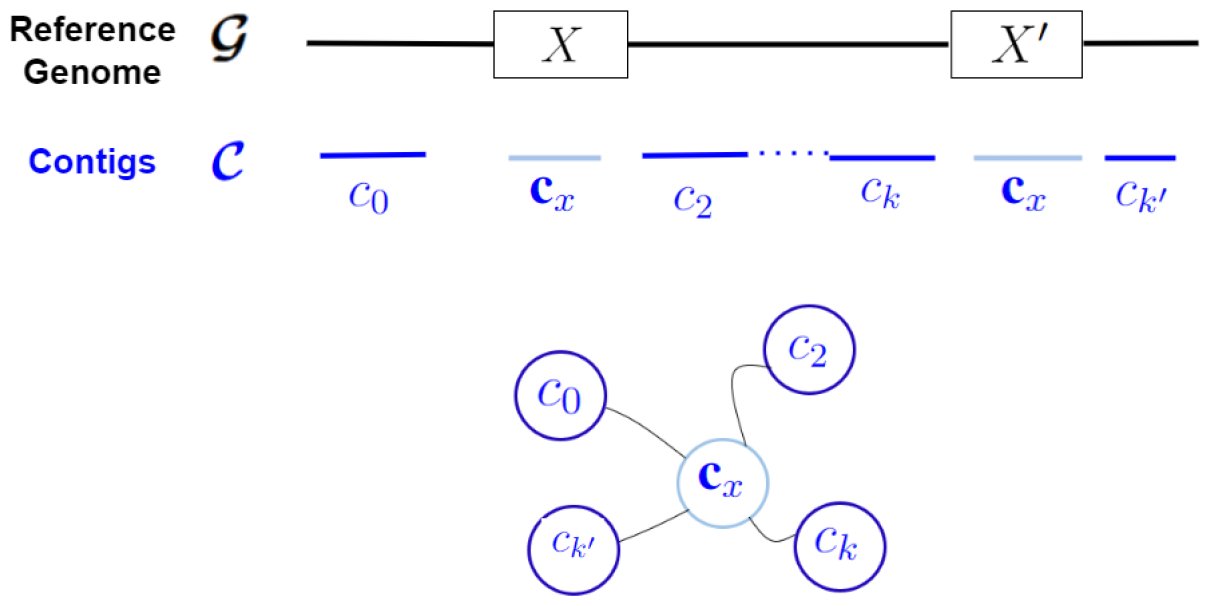
A case of repeats (*X, X*^*′*^) causing cycles branching around contigs.

An important implication of this corollary is that our algorithm is careful in using contigs that fall inside repetitive regions. In other words, if a contig appears as a non-terminal vertex along a path, then its predecessor and successor contigs are those to which this contig exhibits maximum support in terms of its long read based links. While it is not possible to guarantee full correctness, the wiring algorithm uses long read information in order to reduce the chances of repetitive regions causing chimeric scaffolds.

#### Algorithm 1 Wiring Heuristic

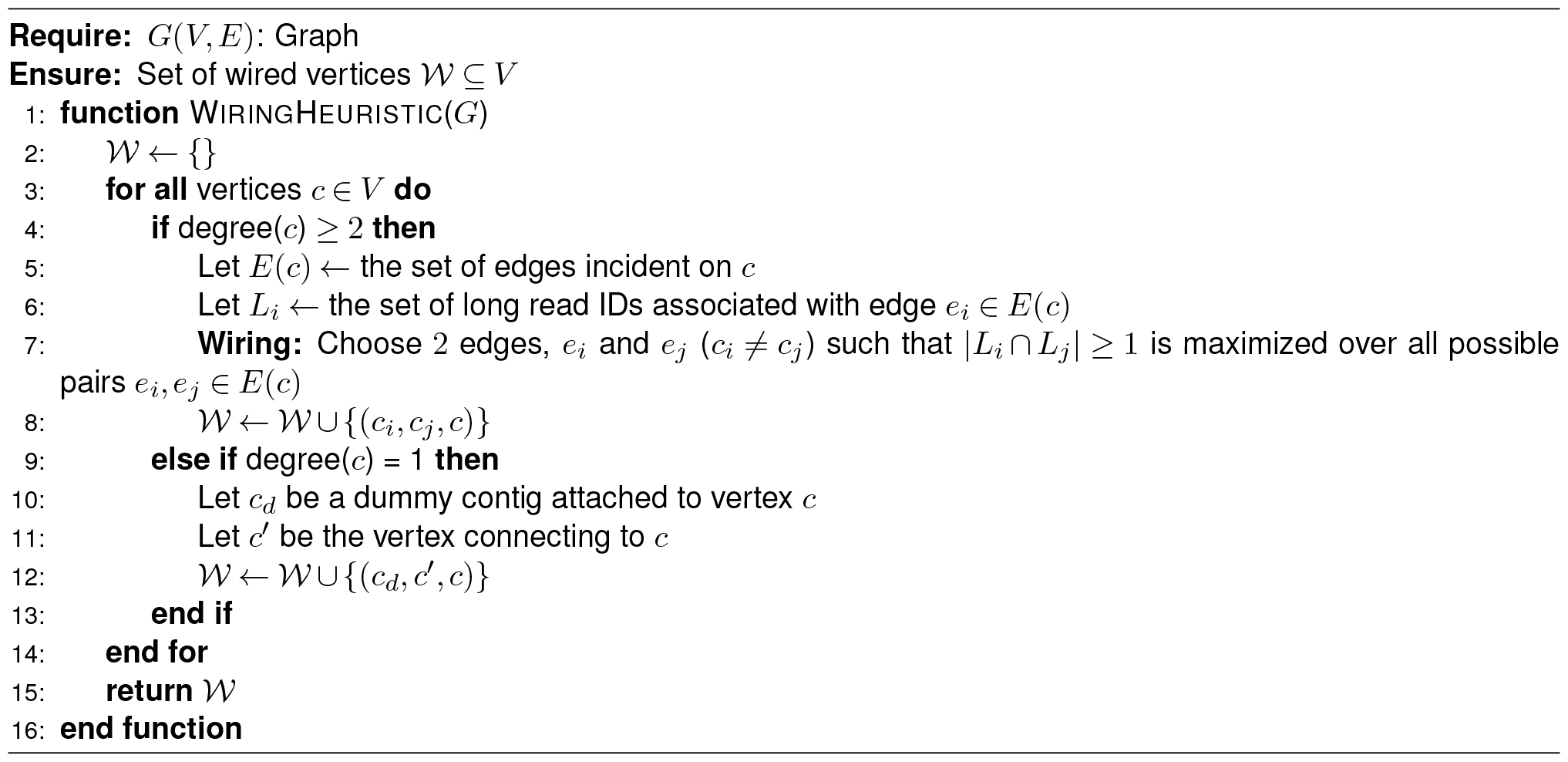

### Step: Parallelized contig batch assembly

As the next step to wiring and path enumeration, we use the paths enumerated to build the output sequence (partial) scaffolds from this phase. To implement this step in a scalable manner, we make a simple observation that the paths enumerated all represent a disjoint set of partial scaffolds. Therefore, we use a partitioning strategy to partition the set of paths into fixed size batches (each containing *s* contigs), so that these independent batches can be fed in a parallel way, into a standalone assembler that can use both the contigs and long reads of a batch to build the sequences corresponding to the partial scaffolds. We refer to this parallel distributed approach as contig batch assembly.

The assembly of each batch is performed in parallel using any standalone assembler of choice. We used `Hifiasm` [5] for all our experiments. By executing contig-long read pairs in parallel batches, this methodology yields one or more scaffolds per batch, contributing to enhanced scalability in assembly processes. Furthermore, the selective utilization of long reads mapped to specific contig batches significantly reduces memory overhead, mitigating the risk of misassemblies that might arise from using the entire long read set which is evident in the results.

This strategy not only reduces memory utilization but also minimizes the potential for misassembly errors that could occur when unrelated sequences are combined.

### Phase: longread island construction

The first phase of contig expansion, only focuses on expanding contigs using long reads that map on either side. This can be thought of a seed-and-extend strategy, where contigs are seeds and extensions happen with the long reads. However, there could be regions of the genome that are not covered by this contig expansion step. Therefore, in this phase, we focus on constructing “longread islands” to cover these gap regions. See Figure 1 for an ilustration of these long read islands. This is achieved in two steps:

a. First we detect all long reads that do not map to any of the first generation partial scaffolds (generated from the contig expansion step). More specifically, we give as input to `JEM-mapper` the set of all unused long reads (i.e., unused in the partial scaffolds) and the set of partial scaffolds output by the contig expansion phase. Any long read that maps to previous partial scaffolds are not considered for this phase. Only those that remain unmapped correspond to long reads that fall in the gap regions between the partial scaffolds.
b. Next, we use the resulting set of unmapped long reads to build partial scaffolds. This is achieved by inputing the unmapped long reads to `Hifiasm`. The output of this phase represent the second generation of partial scaffolds, each corresponding to a long read island.

### Phase: link scaffolds with bridges

In the last phase, we now link the first and second generations of partial scaffolds using any long reads that have been left unused so far. The objective is to bridge these two generations into longer scaffolds if there is sufficient information in the long reads to link them. Note that from an implementation standpoint this is same as for contig expansion, where the union of first and generation partial scaffolds serve as the “contigs” and the rest of the unused long reads serve as the long read set.

### Complexity analysis

Recall that *m* denotes the number of input long reads in *ℒ*, and *n* is the number of input contigs in *𝒞*. Let *p* denote the number of processes used by our parallel program.

Out of the three major phases of `Maptcha`, the `contig expansion` phase is the one that works on the entire input sets (*ℒ* and *𝒞*). The other two phases work on a reduced subset of long reads (unused by the partial scaffolds of the prior scaffolds) and the set of partial scaffolds (which represents a smaller size compared to *𝒞*). For this reason, we focus our complexity analysis on the `contig expansion` phase.

In the `contig expansion` phase we have the following steps:

i. *Mapping long reads to contigs*: `JEM-mapper` [30] is an alignment-free distributed memory parallel implementation and hence processes load the long reads and contigs in a distributed manner. The dominant step is sketching the input sequences (long reads or contigs). Given that the number of long reads is expected to be more than the number of contigs (due to sequencing depth), the complexity can be expressed as 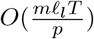, where *𝓁*_*l*_ is average long read length and *T* denotes the number of random trials used within its minhash sketch computation.
ii. *Graph construction*: Let the list of mapped tuples ⟨*c, r*⟩ from the previous step contain *𝒯* tuples. These *𝒯* tuples are used to generate the contig graph by first sorting all the tuples by their long read IDs to aggregate all contigs that map to the same ID. This can be achieved using an integer sort that scans the list of tuples linearly and inserts into a lookup table for all long read IDs—providing a runtime of *O*(*m* +*𝒯*) time. Next, this lookup table is scanned one long read ID at a time, and all contigs in its list are paired with one another to create all the edges corresponding to that long read. The runtime of this step is proportional to the output graph size (*G*(*V, E*)), which contains *n* vertices (one for each contig), and |*E*| is the number of edges corresponding to all contig pairs detected. Our implementation performs this graph construction in a multithreaded mode.
iii. *Wiring heuristic*: For the wiring step, each node detects a pair of edges incident on it that has the maximum intersection in the number of long read IDs. This can be achieved in time proportional to *O*(*d*^2^) where *d* is the average degree of a vertex. The subsequent step of path enumeration traverses each edge at most once. Since both these steps are parallelized, the wiring heuristic can be completed in 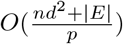 time.
iv. *Contig batch assembly* : The last step is the contig batch assembly, where each of the set of enumerated paths are partitioned into *b* batches, and each batch is individually assembled (using `Hifiasm`). As this step is trivially parallelizable, this step takes 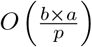 time, where *a* is the average time taken for assembling any batch.

In our results, we show that the `contig expansion` phase dominates the overall runtime of execution (shown later in Figure 6).

**Figure 6.**
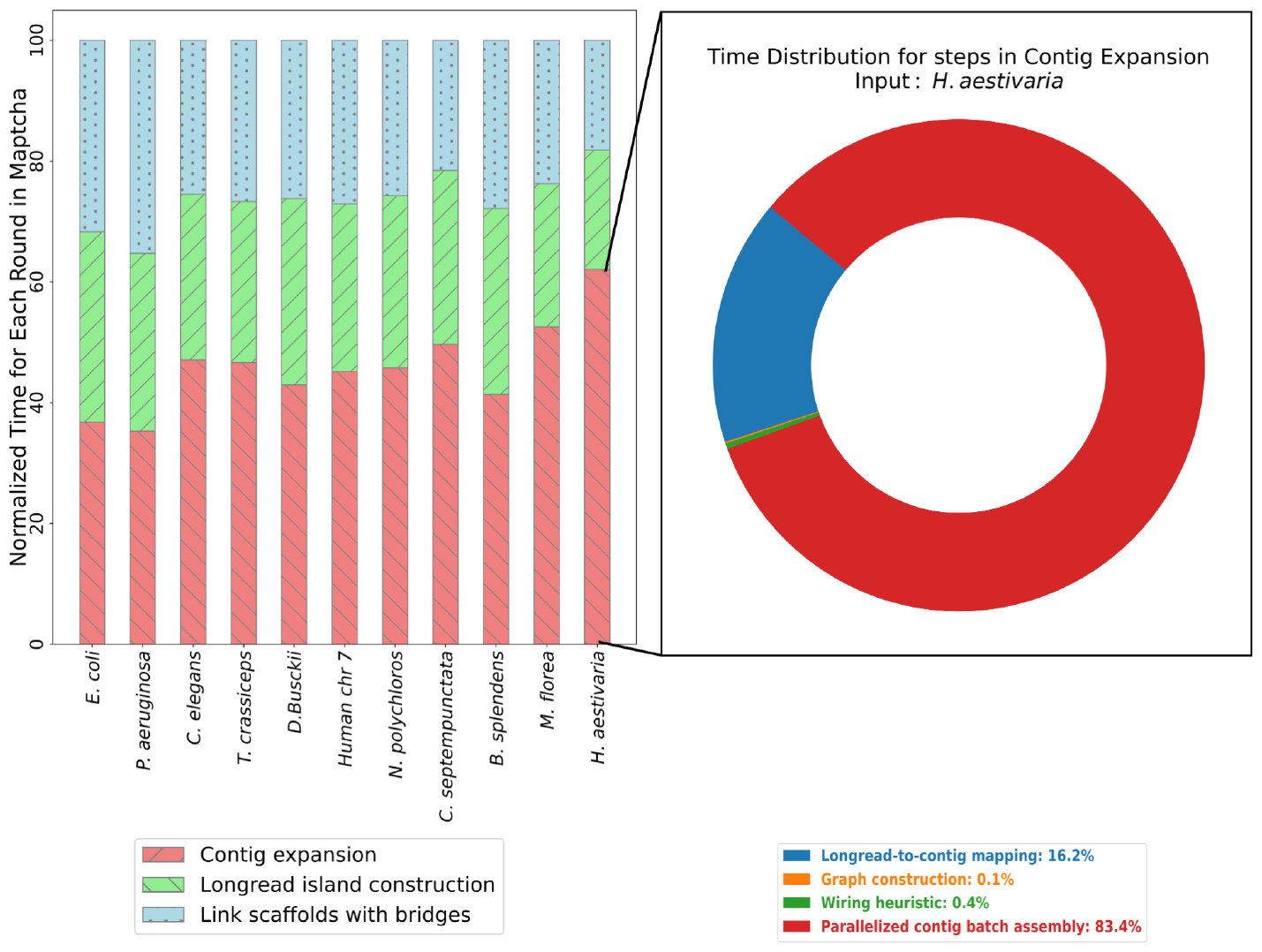
(a) Normalized runtime breakdown for the different rounds of Maptcha pipeline for *p* = 64. (b) Normalized runtime breakdown for different steps in the contig expansion round for input *H. aestivaria*.

The space complexity of `Maptcha` is dominated by the size to store the input sequences and the size of the contig graph—i.e., *O*(*N* + *M* + *n* +|*E*|).

## Results

### Experimental setup

#### Test inputs

For all our experiments, we used a set of input genomes (from various families) downloaded from the NCBI GenBank [1]. These genome data sets are summarized for their key statistics in Table 1. Using the reference for each genome, we generated a set of contigs and a set of long reads as follows. The set of test input contigs (*𝒞*) were generated by first generating and then assembling a set of Illumina short reads using the ART Illumina simulator [17], with 100× coverage and 100bp read length. For short read assembly, we used the Minia [6] assembler. As for the set of test long reads (*ℒ*), we used the Sim-it PacBio HiFi simulator [11], with a 10× coverage and long read median length 10Kbp. Furthermore, note the length divergences in both *𝒞* and *ℒ*.

**Table 1.** Input data sets used in our experiments. All inputs were downloaded from NCBI GenBank [1].

| Input genome | | $\mathcal{C}$: Contig statistics (Minia contigs) | | | $\mathcal{L}$: Long read statistics (HiFi simulated reads) | | |
| --- | --- | --- | --- | --- | --- | --- | --- |
| Genome | Genome length (in bp) | No. contigs ($\geq 1,000$bp) (n) | Total length in bp (N) | Contig length (avg.±std.dev) | No. long reads (m) | Total length in bp (M) | Read length (avg.±std.dev) |
| *E. coli* | 4,641,652 | 330 | 4,499,289 | 13,634 ± 14,139 | 4,541 | 46,312,093 | 10,198 ± 3,420 |
| *P. aeruginosa* | 6,264,404 | 370 | 6,093,817 | 16,469 ± 19,052 | 6,122 | 62,511,066 | 10,210 ± 3,372 |
| *C. elegans* | 100,272,607 | 18,054 | 77,564,568 | 4,296 ± 5,585 | 98,103 | 1,001,075,296 | 10,204 ± 3,396 |
| *T. crassiceps* | 107,053,072 | 4,976 | 90,108,186 | 18,108 ± 31,064 | 104,679 | 1,065,911,598 | 10,182 ± 3,390 |
| *D. busckii* | 118,492,362 | 28,505 | 99,697,088 | 3,497 ± 3,498 | 123,781 | 1,258,903,798 | 10,170 ± 3,406 |
| *Human chr 7* | 159,345,173 | 34,921 | 96,494,010 | 2,763 ± 2,090 | 156,285 | 1,591,064,955 | 10,180 ± 3,390 |
| *N. polychloros* | 398,112,776 | 91,698 | 220,924,212 | 2,409 ± 1,824 | 389,895 | 3,973,622,942 | 10,191 ± 3,398 |
| *C. septempunctata* | 398,868,586 | 57,938 | 136,903,130 | 2,362 ± 2,566 | 390,797 | 3,981,681,897 | 10,188 ± 3,398 |
| *B. splendens* | 441,388,503 | 73,785 | 322,195,214 | 4,366 ± 4,466 | 432,230 | 4,404,143,269 | 10,189 ± 3,393 |
| *M. florea* | 485,103,743 | 86,826 | 218,502,680 | 2,516 ± 1,775 | 474,914 | 4,836,563,328 | 10,184 ± 3,393 |
| *H. aestivaria* | 501,713,186 | 76,767 | 154,096,503 | 2,007 ± 1,343 | 491,533 | 5,005,317,758 | 10,183 ± 3,397 |

#### Qualitative evaluation

To evaluate the quality of of the scaffold outputs produced by `Maptcha`, we used Quast [15] which internally maps the scaffolds against the target reference genome and obtains key qualitative metrics consistent with literature, such as NG50 length, largest alignment length, number of misassemblies, and genome fraction (the percentage of genome recovered by the scaffolded assembly). For a comparative evaluation against a state-of-the-art hybrid scaffolder, we compared the quality as well as runtime performance of `Maptcha` against that of `LRScaf` [29].

#### Test platform

All experiments were conducted on a distributed memory cluster with 7 compute nodes, each with 64 AMD Opteron™ (2.3GHz) cores and 128 GB DRAM. The nodes are interconnected using 10Gbps Ethernet and share 190TB of ZFS storage. The cluster supports OpenMPI and OpenMP. Note that only `Maptcha` is a parallel implementation that can run on distributed memory clusters. All other codes we compared against (`LRScaf`, `Hifiasm (only-LR)`) are multi-threaded and they were run using all 64 threads available on a single node of this cluster.

### Qualitative evaluation

#### Scaffold quality

First, we report on the qualitative evaluation for `Maptcha`, for its hybrid assembly quality. Table 2 shows the quality by the various assembly metrics alongside the quality values for `LRScaf`—for all the inputs tested. The same inputs were provided into both tools. Note that the assembly quality for `Maptcha` shown are for the final output set of scaffolds produced by the framework (i.e, after its `link scaffolds with bridges` phase).

**Table 2.** Quality metrics for our test inputs. Symbol − indicates that these results could not be collected in time on the same system i.e 6 hours.

| Input | Method | NG50 | Largest Alignment (bp) | Genome Coverage % | Missassemblies | Duplication Ratio |
| --- | --- | --- | --- | --- | --- | --- |
| *E. coli* | LRScf | 4,499,158 | 3,541,973 | 97.12 | 0 | 1.03 |
|  | Maptcha | 4,448,034 | 4,509,043 | 99.87 | 0 | 1 |
| *P. aeruginosa* | LRScf | 3,780,771 | 3,703,369 | 97.96 | 1 | 1.03 |
|  | Maptcha | 6,101,601 | 6,102,781 | 98.69 | **0** | 1 |
| *C. elegans* | LRScf | 1,080,091 | 5,019,647 | 83.68 | 46 | 1.23 |
|  | Maptcha | **15,736,218** | **17,718,942** | 99.81 | 11 | **1** |
| *T. crassiceps* | LRScf | 171,559 | 1,108,210 | 86.86 | 501 | 1.02 |
|  | Maptcha | **4,805,993** | **12,352,928** | **98.68** | **21** | **1** |
| *D. busckii* | LRScf | 2,597,298 | 13,199,135 | 91.41 | 42 | 1.11 |
|  | Maptcha | **3,533,287** | 23,381,820 | **95.69** | **0** | 1.01 |
| *Human chr 7* | LRScf | 4,499,158 | 3,541,973 | 97.12 | 0 | 1.03 |
|  | Maptcha | **81,166,983** | **81,144,021** | 99.71 | **2** | **1** |
| *N. polychloros* | LRScf | — | — | — | — | — |
|  | Maptcha | **13,933,406** | **18,337,428** | **99.92** | **15** | **1** |
| *C. septempunctata* | LRScf | — | — | — | — | — |
|  | Maptcha | **24,570,419** | 40,568,023 | **99.95** | 25 | **1** |
| *B. splendens* | LRScf | — | — | — | — | — |
|  | Maptcha | **18,757,076** | **31,409,892** | **99.01** | **54** | **1.1** |
| *M. florea* | LRScf | — | — | — | — | — |
|  | Maptcha | **44,041,601** | **54,515,726** | **97.91** | 67 | **1** |
| *H. aestivaria* | LRScf | — | — | — | — | — |
|  | Maptcha | **18,646,319** | **29,935,333** | **99.83** | 34 | **1** |

We observe from Table 2 that `Maptcha` is able to produce a high quality assembly, reaching nearly 99% genome coverage with high NG50 and largest alignment lengths, low misassembly rate, and a near perfect (1.0) duplication ratio, for all the test inputs. In comparison, `LRScaf`’s quality significantly varies from small to medium range inputs. For the smaller genomes such as *E. coli*, `LRScaf` is able to produce NG50 values that are competitive. However, as the genome lengths grow (e.g., *C. elegans, T. crassiceps*) the assemblies produced are much shorter and fragmented compared to `Maptcha`. For instance, on *T. crassiceps*, the NG50 value for `Maptcha` is about 28× larger compared to that of the value for `LRScaf`. The same trend, with `Maptcha` outperforming `LRScaf`, can be observed for the other assembly metrics including largest alignment length, number of misassemblies, and duplication ratio. Note that it was *not* possible to gather the assembly results for `LRScaf` on the larger genomes (entries shown as −) because the corresponding runs could not complete in less than 6 hours of allotted time on the compute cluster.

We further examined the growth of contigs and incremental improvement in assembly quality through the different scaffolding phases of `Maptcha`. Table 4 shows these results, using NG50 lengths output from these different phases as the basis for this improvement assessment. Figure S4 shows the increase across all three phases in log-scale. The main observations are as follows:

- It can be seen that Minia-assembled contigs for larger genomes initially have NG50 measurements ranging from 1-3 Kbp, but after the `contig expansion` phase, a significant increase in NG50 is observed, often exceeding 200-fold. For instance, inputs such as *C. septempunctata, M. florea*, and *H. aestivaria* show a significant increase of NG50 values from around 2 Kbp for the initial contigs to over 400 Kbp post-`contig expansion` phase. The long reads act as connectors between the shorter contigs, resulting in longer partial scaffolds and explaining the increase in NG50.
- The subsequent phase of `longread island construction` modestly increases NG50 but its main contribution is to provide a more comprehensive genome coverage in regions that are not covered by contigs.
- The final phase of linking partial scaffolds with remaining long reads in `Maptcha` results in a significant surge in NG50 (up to 1,000x for larger genomes). This phase, similar to the `contig expansion` phase, shows the greatest increase in NG50 among all phases. The average length of these partial scaffolds being longer contributes to this increase.

### Performance evaluation

Next, we report on the runtime and memory performance of `Maptcha` and compare that with `LRScaf`. Table 3 shows these comparative results for all inputs tested. All runs with `Maptcha` were obtained by running it on the distributed memory cluster using *p* = 64 processes—more specifically on 4 compute nodes, each running 16 processes. For `LRScaf`, we ran it in its multithreaded mode on 64 threads on a single node of the cluster. Note that in parallel computing, distributed memory systems support larger aggregate memory but at the expense of incurring communication (network) overheads, which do not appear in multithreaded systems running on a single node. However to enable a fair comparison on equivalent number of resources, we tested both on the same number (*p*) of processes, with `Maptcha` running in distributed memory mode while `LRScaf` running on shared memory. For all runs reported for the performance evaluation, we ran `Maptcha` with a batch size of 8,192 in the batch assembly step.

**Table 3.**
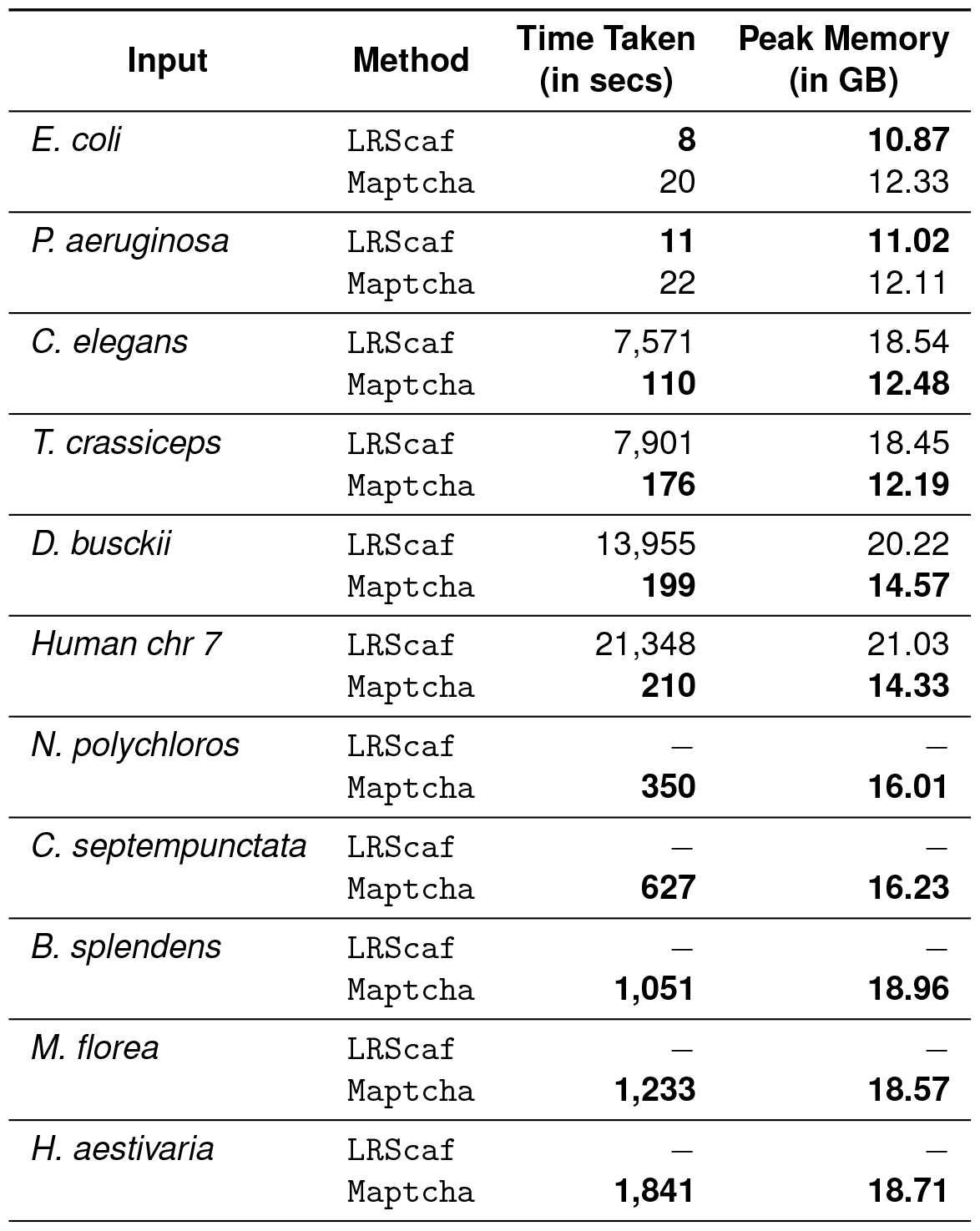
Performance evaluation for our test inputs. Symbol − indicates that these results could not be collected in time on the same system i.e 6 hours.

**Table 4.** The increases in the values of NG50 achieved through the Maptcha phases starting from the input contigs to the final scaffolds.

| Input genome |  | NG50 after each phase of Maptcha (in bp) |  |  |
| --- | --- | --- | --- | --- |
| Genome | Contigs<br>NG50 (in bp) | contig expansion | longread island construction | link scaffolds with bridges |
| *E. coli* | 22,175 | 348,034 | 357,799 | 4,448,034 |
| *P. aeruginosa* | 29,539 | 310,601 | 329,562 | 6,101,601 |
| *C. elegans* | 4,481 | 294,365 | 294,400 | 15,736,218 |
| *T. crassiceps* | 40,139 | 351,834 | 358,177 | 4,805,993 |
| *D. busckii* | 3,678 | 303,316 | 311,207 | 3,533,287 |
| *Human chr 7* | 1,667 | 384,512 | 493,519 | 81,116,983 |
| *N. polychloros* | 1,260 | 325,208 | 474,756 | 13,933,406 |
| *C. septempunctata* | 2,563 | 457,265 | 599,471 | 24,570,419 |
| *B. splendens* | 3,998 | 410,300 | 411,111 | 18,757,076 |
| *M. florea* | 2,740 | 483,063 | 587,311 | 44,041,601 |
| *H. aestivaria* | 2,118 | 439,243 | 547,441 | 18,646,319 |

The results in Table 3 show clearly that `Maptcha`, despite running in distributed memory, clearly outperforms `LRScaf` in its run-time performance. For instance, even on the medium-sized inputs such as *C. elegans*, `Maptcha` is able to complete nearly 70× faster than `LRScaf`, reducing the time to solution from over 2 hours (`LRScaf`) to 110 seconds (`Maptcha`). For the larger inputs, this speedup factor further improves, with `Maptcha` showing over two orders of magnitude faster run-times compared to `LRScaf`. We note that for the 5 largest inputs (out of the 11 test inputs), we were not able to obtain performance results for `LRScaf` since those runs could not complete within the allotted 6 hours limit of the cluster. In contrast, `Maptcha` was able to complete within 20 to 30 minutes on the largest three inputs (*B. splendens, M. florea*, and *H. aestivaria*).

Table 3 also shows the memory used by the two tools for all the inputs. For `Maptcha`, recall that the memory is primarily dictated by the memory needed to produce the batch assemblies (which are partitioned into batches). Due to batching, even though the input genome size is increased, the number of contigs that anchor a batch is kept about the same, ensuring a way to control the memory needed to run large assemblies in a scalable fashion. This is the reason why despite growing input sizes, the peak memory used by `Maptcha` stays approximately steady (under 20 GB).

We also studied the runtime breakdown of `Maptcha` across its different phases. This breakdown is shown normalized for each input in Figure 6a (left), all running on *p* = 64 processes. It can be observed that the `contig expansion` phase is generally the most time consuming phase, occupying anywhere between 40% to 60% of the runtime, with the other two phases roughly evenly sharing the remainder of the runtime. Figure 6b (right) further shows how the run-time is distributed within the `contig expansion` phase. As can be noted, more than 80% of the time is spent in the batch assembly step, while the remainder of the run-time is spent mostly on mapping.

### Effect of batch size on NG50 and run-time

Figure 7 shows the impact of varying batch sizes on NG50 and processing time, using the *H. aestivaria* genome as an example. Recall that the batch size is the number of contigs that are used to anchor each batch along with their respective long reads that map to those contigs. Subsequently, each batch is provided to a standalone assembly (using `Hifiasm`) to produce the assemblies for the final scaffolds. We experimented with a wide range of batch size, starting from 32, and until 16K. As anticipated, smaller batch sizes exhibit reduced processing times due to the smaller assembly workload per batch. However, if a batch is too small then the resulting assembly quality is highly fragmented (resulting in small NG50 values) as can be observed. Conversely, larger batch sizes necessitate longer processing times (e.g., batch size 32 requiring approximately 280 seconds, while 8K batch size requires 1,841 seconds). But the NG50 metric significantly improves—e.g., NG50 size improvement from 93Kbp to 1.8Mbp from a batch size of 32 to 8K.

**Figure 7.**
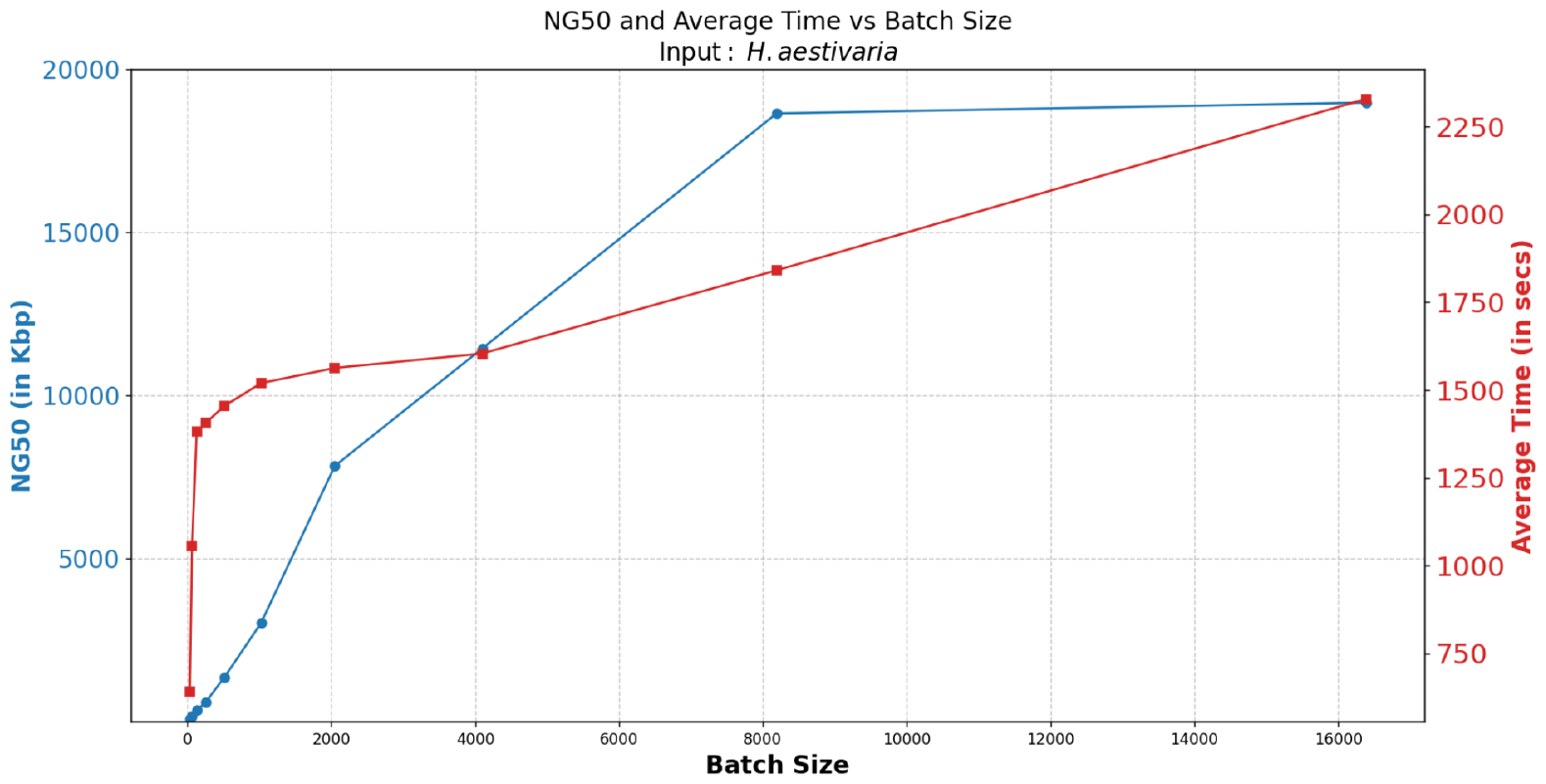
Effect of batch size on NG50 and average time taken for input *H. aestivaria*.

We found that increasing the batch size from 8K to 16K resulted in a slight increase in NG50 (1.86Mbp to 1.89Mbp), but also a significant increase in processing time (1,841 seconds to 2,329 seconds). Since the increase in NG50 was not substantial enough to justify the longer processing time, we decided to use the batch size of 8K for all our tests.

#### Coverage experiment with Maptcha (hybrid) and Hifiasm (only-LR)

One of the main features of a hybrid scaffolding workflow is that it has the potential to build incrementally on prior constructed assemblies using newly sequenced long reads. This raises two questions: a) how does the quality of a hybrid workflow compare to a standalone long read-only workflow? b) can the information in contigs (or prior constructed assemblies) be used to offset for lower coverage sequencing depth in long reads?

To answer these two questions, we compared the `Maptcha` scaffolds to an assembly produced directly by running a standalone long read assembler but just using the long reads. For the latter, we used `Hifiasm` and denote the corresponding runs with the label `Hifiasm (only-LR)` (to distinguish it from the hybrid configuration in `Maptcha`). Analysis was performed using different coverages (1x, 2x, 3x, 4x, 8x, and 10x) for the long read data set, for the *H. aestivaria* input, and focusing on performance metrics of NG50, execution time, and peak memory utilization.

The results shown in Table 5 for this experiment, revealed that at lower coverages (1x and 2x), `Hifiasm (only-LR)` and `Maptcha` demonstrated relatively comparable performance. However, as the long read coverage increased, `Maptcha` exhibited better NG50 quality over `Hifiasm (only-LR)`, demonstrating the value of adding the contigs in growing the scaffold length. For instance, at 4x coverage, `Maptcha` yielded a substantially longer NG50 (ten-fold increase). The assembly quality becomes comparable for higher coverage settings. These results demonstrate that the addition of prior constructed assemblies can increase the scaffold length compared to long read-only assemblies. However, this value in growing the scaffold length tends to diminish for higher coverage settings—showing that the addition of contigs can be used to offset reduced coverage settings.

**Table 5.**
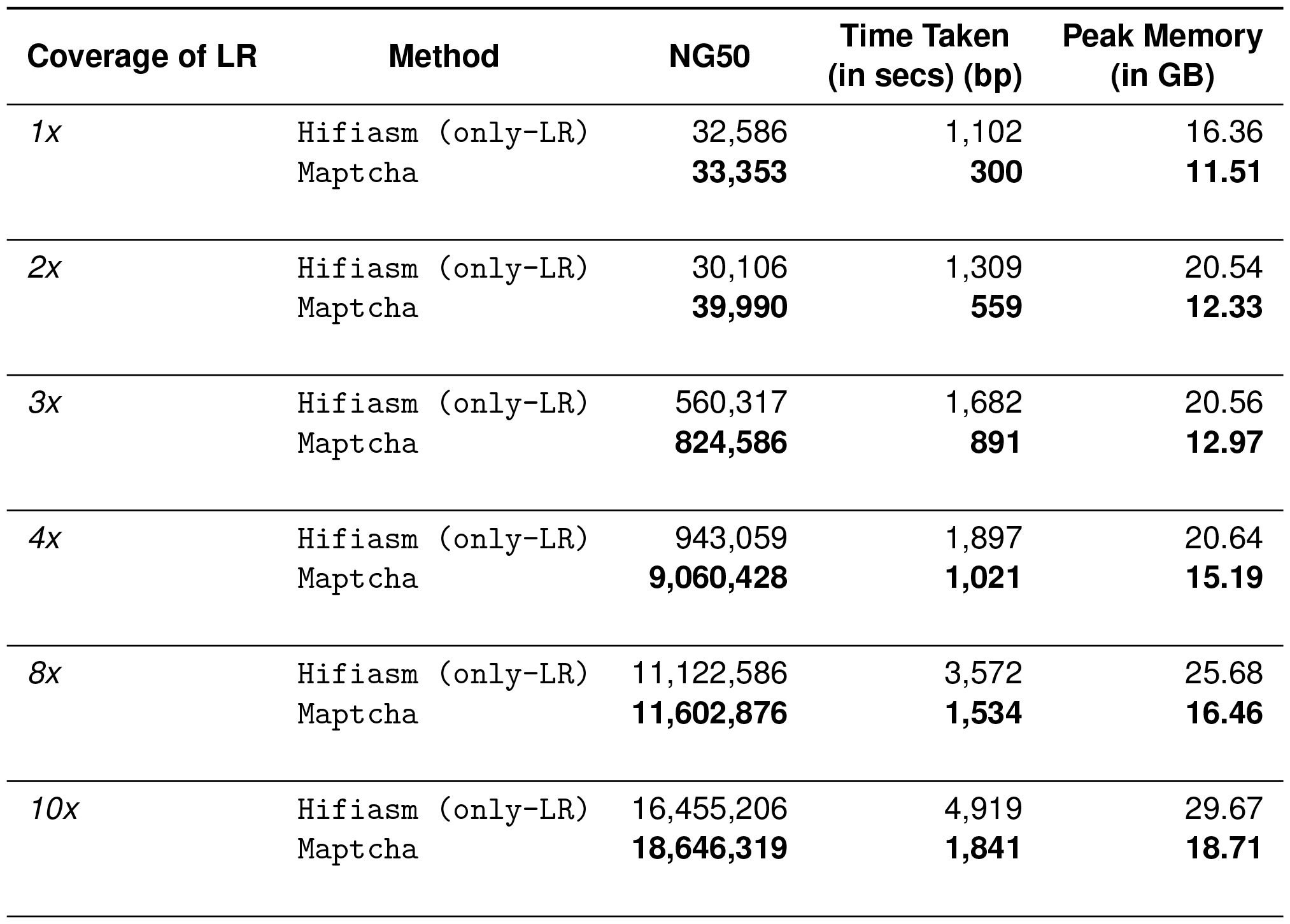
Quality and performance evaluation of running Hifiasm (only-LR) and Maptcha with different coverages of longread on input *H. aestivaria*.

| Coverage of LR | Method | NG50 | Time Taken<br>(in secs) (bp) | Peak Memory<br>(in GB) |
| --- | --- | --- | --- | --- |
| 1x | Hifiasm (only-LR) | 32,586 | 1,102 | 16.36 |
|  | Maptcha | **33,353** | **300** | **11.51** |
| 2x | Hifiasm (only-LR) | 30,106 | 1,309 | 20.54 |
|  | Maptcha | **39,990** | **559** | **12.33** |
| 3x | Hifiasm (only-LR) | 560,317 | 1,682 | 20.56 |
|  | Maptcha | **824,586** | **891** | **12.97** |
| 4x | Hifiasm (only-LR) | 943,059 | 1,897 | 20.64 |
|  | Maptcha | **9,060,428** | **1,021** | **15.19** |
| 8x | Hifiasm (only-LR) | 11,122,586 | 3,572 | 25.68 |
|  | Maptcha | **11,602,876** | **1,534** | **16.46** |
| 10x | Hifiasm (only-LR) | 16,455,206 | 4,919 | 29.67 |
|  | Maptcha | **18,646,319** | **1,841** | **18.71** |

Table 5 also shows that a run-time and memory advantage of `Maptcha` over `Hifiasm (only-LR)`. For instance, `Maptcha` was generally between two and four times faster than `Hifiasm (only-LR)` (e.g., on the 10x input, `Maptcha` takes 1,841 seconds compared to 4,919 seconds taken by `Hifiasm (only-LR)`). Note that internally, `Maptcha` also is using the standalone version of `Hifiasm` to compute its final assembly product. These results show that the `Maptcha` approach of enumerating paths to generate partial scaffolds and distributing those into batches, reduces the overall assembly workload for the final assembly step, without compromising on the quality.

## Conclusions

Genome assembly remains a challenging task, particularly in resolving repetitive regions within a comparable time. In this study, we present `Maptcha`, a novel hybrid scaffolding pipeline designed to combine previously constructed assemblies with newly sequenced high fidelity long reads. As demonstrated, the `Maptcha` framework is able to increase the scaffold lengths significantly, with the NG50 lengths growing by more than four orders of magnitude relative to the initial input contigs. This represents a significant improvement in genomic reconstruction that comes without any compromise in the accuracy of the genome. Furthermore, our method is able to highlight the value added by prior constructed genome assemblies toward potentially reducing the required coverage depth for downstream long read sequencing. In terms of performance, the `Maptcha` software is a parallel implementation that is able to take advantage of distributed memory machines to reduce time-to-solution of scaffolding. The software is available as open source for testing and application at https://github.com/Oieswarya/Maptcha.git.

## Supplementary Material

### Benchmark construction for evaluating the mapping step

In our investigation of mapping quality against contemporary benchmarks, we utilized the `Minimap2` tool, a widely recognized tool for read-to-reference mapping, extensively employed in recent hybrid scaffolding methodologies such as `LRScaf` [29].

For each input in our test data set, we constructed a benchmark to evaluate the longread-to-contig mapping in our scaffolding workflow. These benchmarks were constructed using their mapping information on the reference genome *G*. Figure S1 shows an illustrative example, involving 6 longreads and 7 contigs. More specifically, to determine the ⟨start,end⟩ benchmark coordinates of each contig and of each longread, we mapped the set of contigs and the set of longreads to the reference genome using `Minimap2` [22]. After this mapping, we say a longread *r* ∈ *ℒ maps* to a contig *c* ∈ *𝒞* if and only if their respective coordinates intersect in at least one position of the reference genome. Using this information, we constructed a set of benchmark (for each input): the mapping benchmark consists of tuples of the form ⟨contig, long read ID⟩;

To compare a test output to the corresponding benchmark, we compute the following measures: a) *True Positive (TP):* when a test output matches with the benchmark; b) *False Positive (FP):* when a test output is *not* in the benchmark; and c) *False Negative (FN):* when a benchmark output is *not* in the test output. Consequently, precision is given by 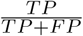, and recall by 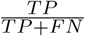.

**Figure S1.**
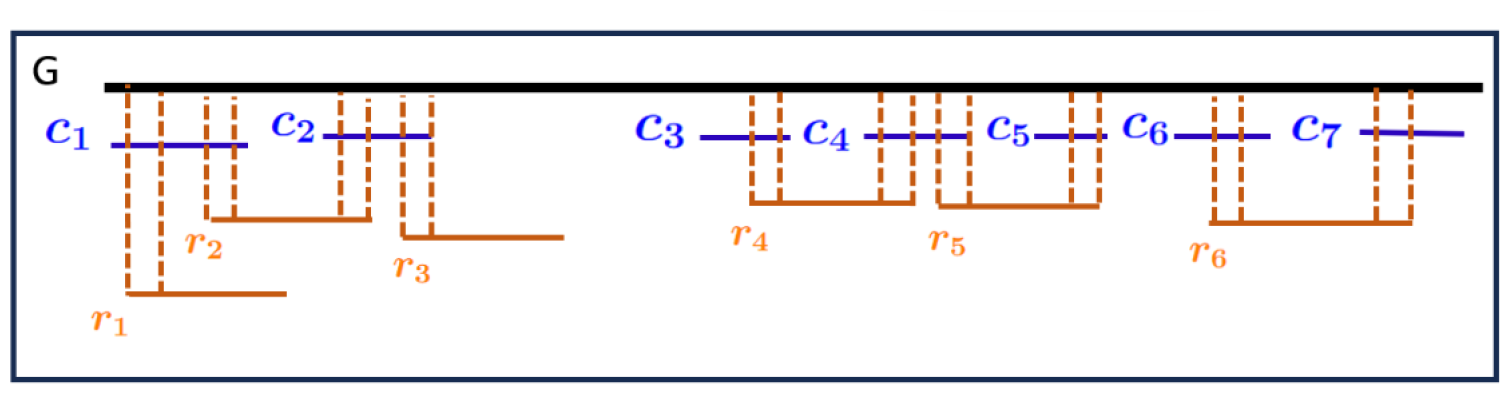
An illustrative example for constructing benchmarks for evaluation of the mapping, using 7 contigs and 6 longreads. The mapping benchmark for this example will contain tuples {⟨*c*_1_, *r*_1_⟩, ⟨*c*_1_, *r*_2_⟩, ⟨*c*_2_, *r*_2_⟩, ⟨*c*_2_, *r*_3_⟩, ⟨*c*_3_, *r*_4_⟩, ⟨*c*_4_, *r*_4_⟩, ⟨*c*_4_, *r*_5_⟩, ⟨*c*_5_, *r*_5_⟩⟨*c*_6_, *r*_6_⟩⟨*c*_7_, *r*_6_⟩}. Consequently, this will contribute to the following edges to the contig graph benchmark: {⟨*c*_1_, *c*_2_⟩, ⟨*c*_3_, *c*_4_⟩, ⟨*c*_4_, *c*_5_⟩, ⟨*c*_6_, *c*_7_⟩}.

#### Mapping quality

The mapping quality of `JEM-mapper` alongside the quality achieved by `Minimap2` is shown in Figure S2. We can observe that the recall achieved by `JEM-mapper` mapping is significantly higher than using `Minimap2`. In all the cases it is well more than the recall rate achieved by `Minimap2`. As for precision, `Minimap2` is better for some inputs like the smaller genomes, *E. coli* and *P. aeruginosa* but `JEM-mapper` is comparable in all cases and higher for some bigger inputs like *C. elegans* and *D. busckii*. Collectively, these results suggest that the MinHash-based scheme holds promise for application in erroneous long read use-cases. The randomness in the procedure to pick the sketches improves recall. This also ensures that there will be substantial contig linking information passed on to the scaffolding step. As for `Minimap2`, the loss in recall can be attributed to the length divergences of the longreads. In particular, we observed that `Minimap2` fails when the length of the contig grows comparable or larger to the longread length; while the stochastic nature of selecting short sketches for `JEM-mapper` coupled with our choice of using only end segments from the longreads, is able to yield better recall rate across the longread length distribution.

**Figure S2.**
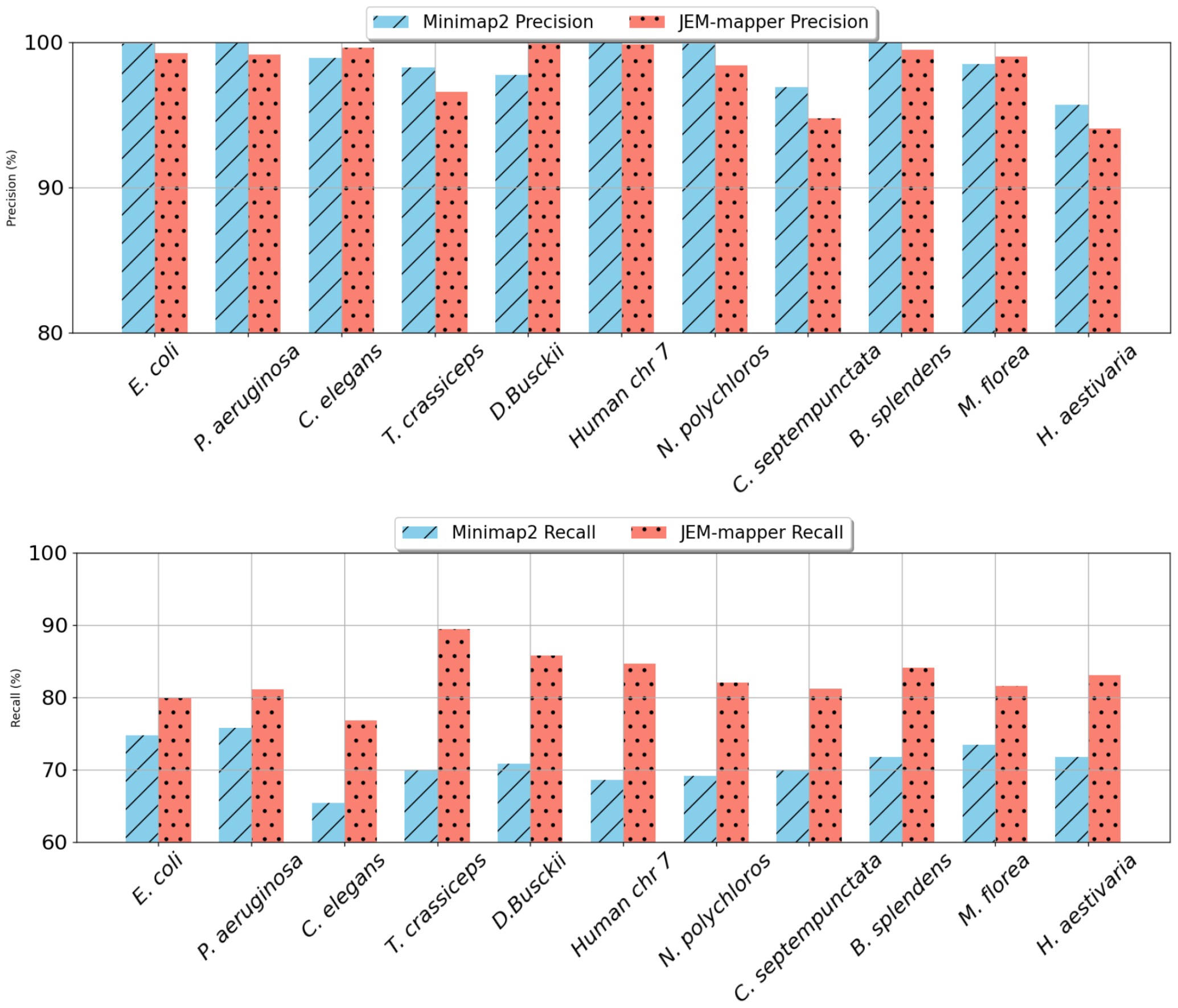
Precision and recall values for the mapping quality comparison between Minimap2 and JEM-mapper.

**Figure S3.**
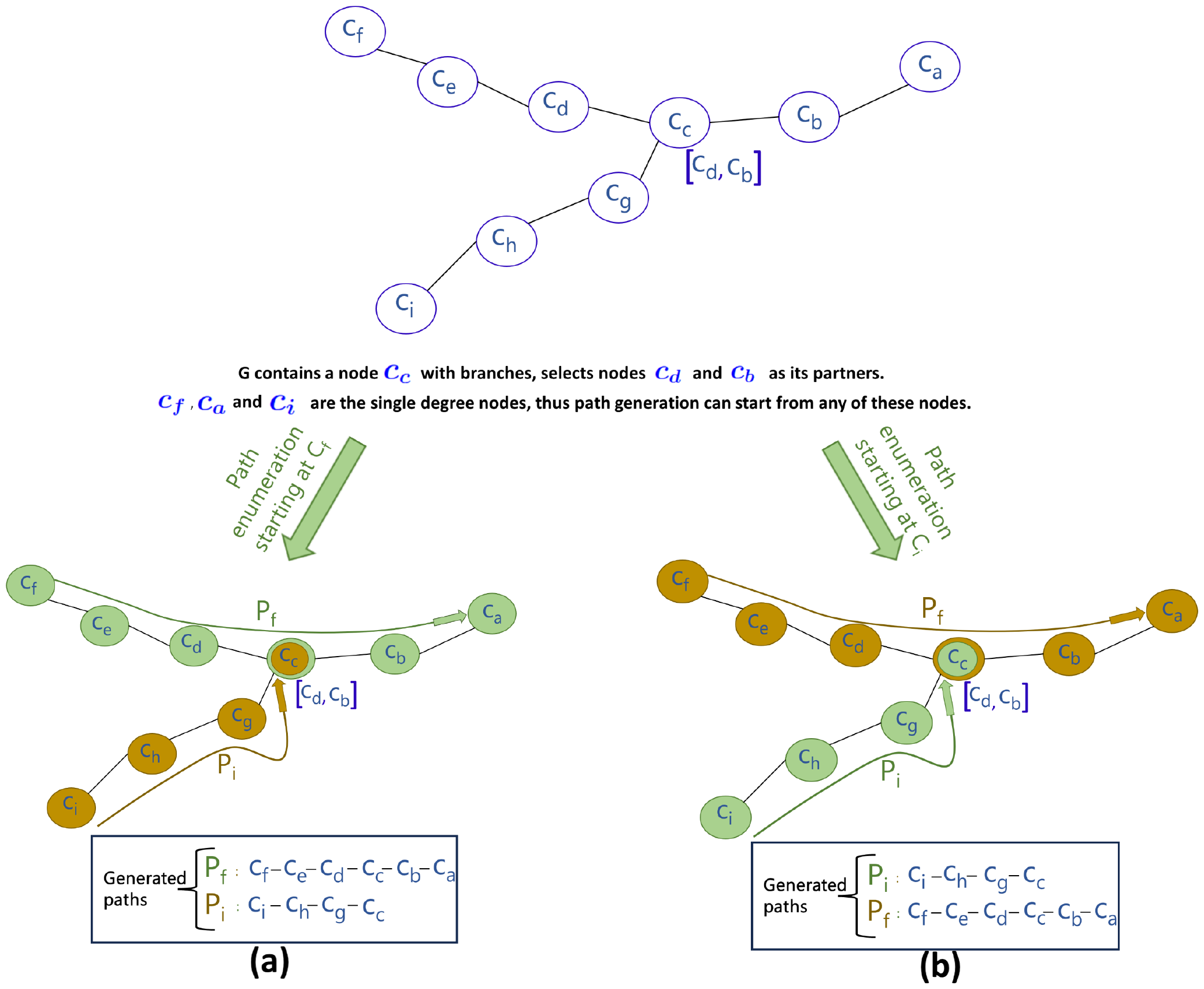
An illustrative example demonstrating the deterministic routing regardless of the traversal origin. This diagram illustrates a scenario where a network node (*c*_*c*_) with branching options (*c*_*d*_ and *c*_*b*_) generates two distinct paths (*P*_*f*_ and *P*_*i*_) from different single-degree node origins (*c*_*f*_ and *c*_*i*_, respectively). Path *P*_*f*_, originating from node *c*_*f*_, traverses through nodes *c*_*e*_, *c*_*d*_, *c*_*c*_, *c*_*b*_, and *c*_*a*_, while path *P*_*i*_, initiated from node *c*_*i*_, progresses through nodes *c*_*h*_, *c*_*g*_ and *c*_*c*_. Despite different starting points, the same two paths are output, showing the deterministic routing property of the path enumeration algorithm.

**Figure S4.**
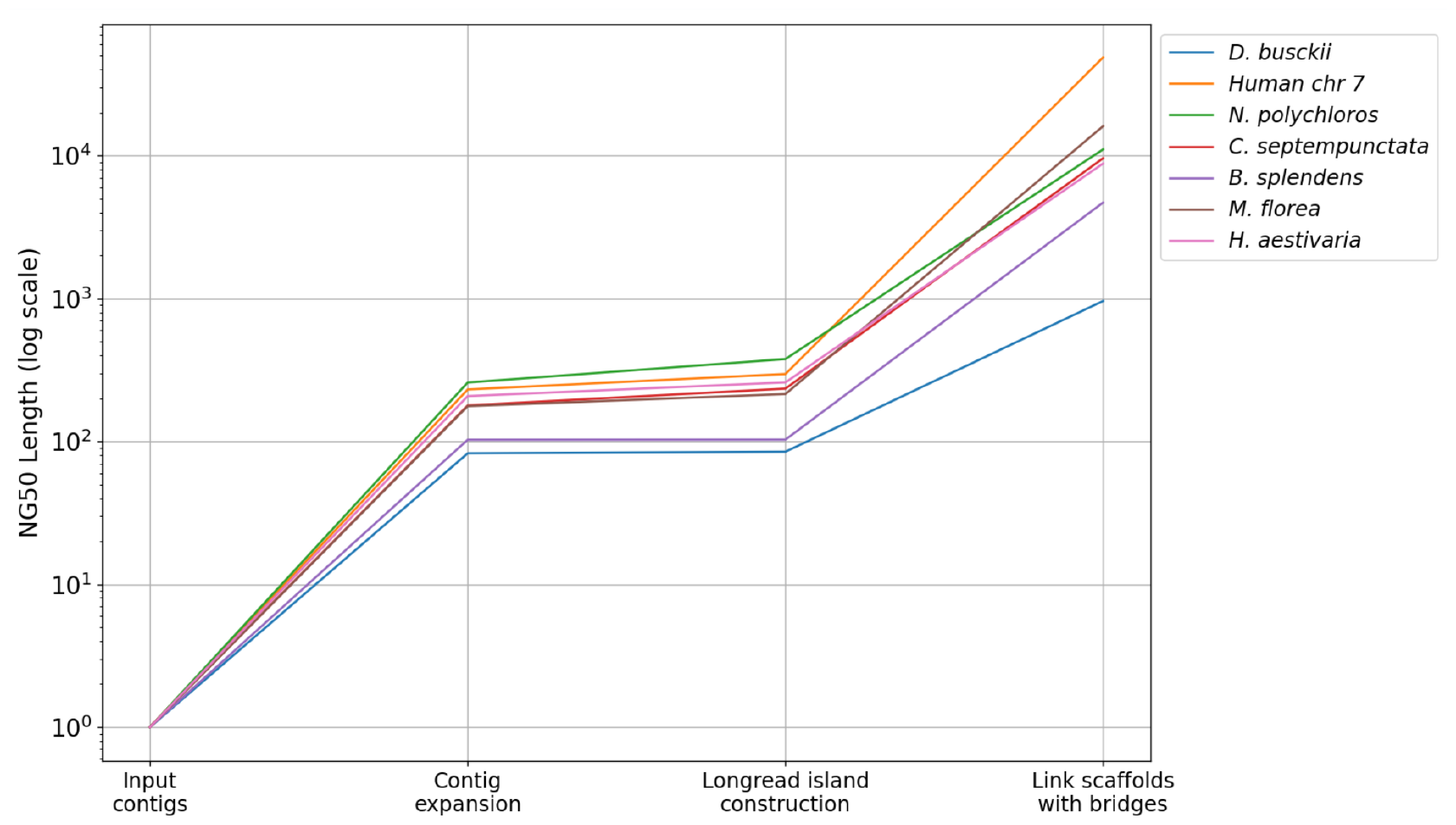
Qualitative improvement of the partial scaffolds (measured in the NG50 lengths) over the different phases of Maptcha. All values (y-axis) are normalized treating the initial contig NG50 length as 1.0 for the respective inputs.

## Footnotes

1 A vertex is said to be *non-terminal* along a path if it appears neither at the start nor the end of that path.

## Notes

### Competing Interest Statement

The authors have declared no competing interest.

